# SARS-CoV-2 S, M and E Structural Proteins Down-modulate HIV-1 LTR Activity and Modulate Endoplasmic Reticulum Stress Responses

**DOI:** 10.1101/2024.07.23.604777

**Authors:** Wejdan Albalawi, Jordan Thomas, Farah Mughal, Kelly J. Roper, Abdullateef Alshehri, Aurelia Kotsiri, Georgios Pollakis, William A. Paxton

**Affiliations:** University of Liverpool, Department of Clinical Infection, Microbiology and Immunology (CIMI), Institute of Infection, Veterinary and Ecological Sciences (IVES), Liverpool, L69 7BE, UK; Department Clinical Laboratory Sciences, College of Applied Medical Sciences, Sakakah 72388, the University of Aljouf, Saudi Arabia; Department of Medical Microbiology, College of Applied Medical Sciences, Najran University, Najran, Saudi Arabia

## Abstract

We have previously shown that the Hepatitis C Virus (HCV) E1E2 envelope glycoprotein can down-modulate HIV-1 long-terminal repeat (LTR) activity through disruption to NF-κB activation. This response is associated with up-regulation of the endoplasmic reticulum (ER) stress response pathway. Here we demonstrate that the SARS-CoV-2 S, M and E but not the N structural protein can perform similar down-modulation of HIV-1 LTR activation and in a dose-dependent manner in both HEK293 and lung BEAS-2B cell-lines and interpreted as a result of NF-κB down-modulation. The effect is highest with the SARS-CoV-2 Wuhan S strain and decreases over-time for the subsequent emerging variants of concern (VOC) with omicron providing the weakest effect. We developed pseudo-typed viral particle (PVP) molecular viral tools that allowed for the generation of cell-lines constitutively expressing separately the four SARS-CoV-2 structural proteins and utilising the VSV-g envelope protein to deliver the integrated gene construct. Differential gene expression analysis (DGEA) was performed on cells expressing S, E, M or N to determine cell activation status. It it was determined that gene expression differences were found in a number of interferon-stimulated genes (ISGs), including IF16, IFIT1, IFIT2 and ISG15 as well as for a number of heat shock protein (HSP) genes, including HSPH1, HSPA6 and HSPBP1 with all four SARS-CoV-2 structural proteins. There were also differences observed with expression patterns of transcription factors with both SP1 and MAVS upregulated in the presence of S, M and E but not the N protein. Collectively the results indicate that gene expression patterns associating with ER stress pathways can be identified with SARS-CoV-2 envelope glycoprotein expression. The results suggest the SARS-CoV-2 can modulate activation of an array of cell pathways resulting in disruption to NF-κB signalling hence providing alterations to multiple physiological responses of SARS-CoV-2 infected cells.

## INTRODUCTION

Since the outbreak of SARS-CoV-2, extensive research has concentrated on understanding the molecular intricacies of viral infection and its impact on cellular biology. The structural proteins of SARS-CoV-2, including spike (S), membrane (M), envelope (E), and nucleocapsid (N) ^[1]^, have become focal points of investigation due to their critical roles in viral replication, assembly, and modulation of host immune responses. The S protein is pivotal for viral entry into host cells ^[2]^, binding to the human ACE2 receptor and serving as a primary target for neutralising antibodies ^[3–5]^. Positive evolutionary selection has shaped the S protein, resulting in the emergence of SARS-CoV-2 variants with enhanced fitness ^[6]^. Understanding how these mutations in the S protein impact viral function is crucial. As of March 2022, the WHO has identified five SARS-CoV-2 strains as variants of concern (VOC): Alpha, Gamma, Delta, and Omicron (WHO, 2022). Better understanding the biological effects of alterations in the S protein aides in deciphering the implications of these variants, guiding ongoing research and public health efforts in managing the evolving landscape of the COVID-19 pandemic.

SARS-CoV-2 S, E and M are structural proteins synthesised, inserted and folded in the ER before being transported to the ER-Golgi apparatus intermediate component (ERGIC), where virus particles are assembled, whereas N protein is translated in the cytoplasm ^[7–10]^. Normal ER function is required for protein synthesis, folding, modification and transport, ER stress is caused by disruption in the structure and function of the ER due to the accumulation of unfold/misfolded proteins ^[11,12]^. The unfolded protein response (UPR) has an emerging function that extends to various cellular processes, including acting in the induction or maintenance of immune responses ^[13]^.

Several studies have consistently observed elevated serum levels of pro-inflammatory cytokines, including interleukin-1 (IL-1), IL-6, and tumour necrosis factor (TNF), in patients with severe COVID-19 symptoms ^[14–16]^. This excessive production of pro-inflammatory cytokines, commonly referred to as cytokine storm, has been associated with acute respiratory distress syndrome (ARDS) and increased mortality in COVID-19 patients ^[17]^. Similar dysregulated immune responses characterised by hyper-production of pro-inflammatory cytokines have been observed in other viral infections such as H5N1 influenza, SARS-CoV, hantavirus pulmonary syndrome, and possibly in HIV-1 infection, where the absence of negative feedback on the immune response contributes to severe disease outcomes ^[18–21]^.

The NF-κB signalling pathway, crucial for coordinating the expression of pro-inflammatory cytokines, consists of five key proteins: p65 (RelA), RelB, c-Rel, p105/p50 (NF-κB1), and p100/52 (NF-κB2). These proteins assemble into unique homo- and heterodimeric complexes, which are transcriptionally active due to their interactions ^[22, 23]^. This pathway plays a critical role in regulating the immune response, particularly during viral infections ^[24]^. The NF-κB pathway can be activated by various stimuli, such as cytokine receptor ligands, pattern-recognition receptors (PRRs), the TNF receptor (TNFR) superfamily, and T-cell and B-cell receptors ^[25]^. The activation of NF-κB occurs through inducible degradation of IκBα, facilitated by a multi-subunit IκB kinase (IKK) complex. Cytokines, growth factors, mitogens, microbes, and infectious agents can all trigger IKK activation ^[26]^. NF-κB, in turn, promotes the expression of numerous pro-inflammatory cytokine genes, leading to further autocrine activation of the NF-κB signalling pathway. Viruses can exploit the NF-κB pathway for their benefit ^[27]^. In the context of COVID-19, SARS-CoV-2 activates NF-κB, leading to pro-inflammatory cytokine production, which correlates with COVID-19 severity ^[28]^. However, the precise mechanisms by which viral modifications influence NF-κB function remain uncertain.

This study aimed to explore how SARS-CoV-2 structural proteins impact NF-κB activity, a key regulator in cellular processes. Using in vitro molecular techniques, we extensively examined the influence of these viral proteins on NF-κB activation. We utilised the HIV-1 LTR promoter to measure NF-κB activation in response to viral proteins. It is imperative to emphasise that HIV-1 replication is predominantly regulated by gene expression originating from the long terminal repeat (LTR), a critical genomic component highly influenced by an array of host factors ^[29]^. The LTR of the viral genome relies on both basal cellular transcription factors and inducible factors, particularly the NF-κB family, along with various other host cellular proteins, for its functioning and regulation ^[30, 31]^. NF-κB is crucial in regulating gene expression across cell types, impacting immune responses, inflammation, and cell survival ^[32, 33]^. Studying the interaction between viral proteins and NF-κB aimed to gain a deeper understanding of the molecular mechanisms underlying the modulation of NF-κB by SARS CoV-2 infection. To investigate this, Oxford Nanopore Technology (ONT) sequencing was used to analyse and determine the impact of SARS-CoV-2 Env glycoprotein expression on the transcriptome of the cell. This investigation aims to uncover the underlying mechanisms by which these viral proteins influence gene expression and explore their impact on transcription factors. Additionally, we aim to analyse the pathways affected by this protein induced changes in gene expression. This analysis will provide valuable insights into the intricate molecular interactions between SARS-CoV-2 proteins and the host cell machinery, contributing to our understanding of SARS-CoV-2 infection pathogenesis and potentially revealing novel therapeutic targets.

## MATERIALS AND METHODS

### Cell Culture

The study used the following human cell lines: the human transfection-competent embryonic kidney cell line, HEK293T, and the human bronchial epithelial cell line, BEAS-2B. All cells were cultured in Dulbecco’s modified eagle medium (DMEM), supplemented with 10% heat-inactivated fetal bovine serum (ΔFBS), 2 mM/ml L-glutamine, and 1% Penicillin-Streptomycin (Pen-Strep). Incubation was carried out at 37°C with 5% CO_2_.

### Plasmid Preparations

Plasmid preparations were conducted through heat shock transformation, followed by DNA purification using Qiagen maxiprep kits. Initially, 2µl of plasmid were added to One Shot TOP10 chemically competent *Escherichia coli*, followed by a 30 min incubation at 42°C and 30 s on ice. After adding 0.5 ml SOC media, the mixture was incubated at 180 rpm for 1 h at 37°C. The culture was plated on antibiotic-selective agar plates, colonies were selected, grown in Brain Heart Infusion Broth (BHI) with ampicillin (100 µg/ml) for miniprep isolation, and subsequently used for maxiprep extraction.

### Transfection of Cell Lines with LTR and SARS-CoV-2 Protein Plasmid Constructs

The SARS-CoV-2 (Wuhan-Hu-1) S, M, E, and N genes were cloned into the pCDNA3.1 expression plasmid (produced by GeneArt Gene Synthesis). This plasmid was also utilised for cloning SARS-CoV-2 variants (produced by GeneArt Gene Synthesis). HEK293T and BEAS-2B cell lines were seeded in 96-well plates at a density of 1.5×10^4^ cells per well and incubated overnight. The medium was then replaced with 50 µl of Opti-MEM. Transfection mixes were prepared by combining 1 ng of Tat plasmid, 6 ng of LTR-luc plasmid, and 12, 6, or 1 ng of SARS-CoV-2 S, M, E and N SARS-CoV-2 variants ^[34]^, or pCDNA plasmid with 2 µl of Opti-MEM. An additional 2 µl of PEI (0.14 μg/μl) was added, and the mix was incubated at room temperature for 30 min. The resulting transfection mixture was added to the cells, which were then incubated for 6 h, followed by a 48-hour incubation in DMEM. Luciferase activity measurement was conducted after this period.

### Stable cell line production

The SARS-CoV-2 (Wuhan-Hu-1) S, M, E, and N genes were integrated into a lentiviral expression plasmid (pGenlenti vector) from Genscript, UK. Lentivirus particles were generated using pseudo-typed viral particles (PVP) with vesicular stomatitis glycoprotein (VSV-g) for enhanced stability and broader tropism. A transient three-plasmid transfection method was employed, with 3×10^6^ HEK293T (lentiX) cells seeded one day before transfection. The lentiviral construct (4500 ng), lentiviral backbone (p8.91, 3000 ng), and VSGg plasmid (2700 ng) ^[35]^ were co-transfected using PEI in OptiMEM. After a 48-hour transfection, lentivirus particles were harvested, filtered (0.45µM) and stored at -80°C. Antibiotic selection was optimised using a puromycin kill-curve on HEK293T cells (1×10^6^). Cells were seeded in two 10 cm^2^ plates in complete DMEM media. After polybrene treatment (8 μg/ml) for 2 minutes, one plate received lentiviral supernatant, while the other served as a control. After two rounds of transduction and a 48-h puromycin treatment (1 μg/ml), media/antibiotic were refreshed every two days until all control cells were dead. Western blot analysis assessed the expression of the gene of interest.

### Flow Cytometry for Transduction Validation

To confirm SARS-CoV-2 protein expression in transduced cells, lentiviral vectors expressing the GFP reporter protein were generated. Lentivirus vector production, infection, and antibiotic selection followed previous protocols. Approximately 1×10^5^ cells were used for GFP signal quantification. After pelleting and washing with FACS buffer, cells were resuspended in 200 μl FACS buffer and analysed using flow cytometry. Flow cytometry data were analysed with FlowJo version 7.10.0.

### Validation of SARS-CoV-2 Stable Cells

Western blotting was employed to assess the expression levels of SARS-CoV-2 S, N, E and N proteins in the transduced cells. Cells (1×10^6^) were washed with ice-cold dPBS and lysed using radioimmunoprecipitation assays (RIPA) buffer supplemented with a 1% protease inhibitor cocktail. The protein concentration was determined using the Bradford assay with the Protein Assay kit. Subsequently, 50 µg of cell lysates were mixed with 4x NuPage LDS Sample Buffer and 10x NuPage reducing agent, incubated at 72°C for 10 min, and separated using polyacrylamide gels (NuPAGE 12% Bis-Tris Gels) via electrophoresis at 120 V for 60 min. The separated proteins were transferred to iBlot™ 2 PVDF Mini Stacks membranes using the iBlot 2 Dry Blotting system. The membranes were blocked using the iBind solution kit and incubated with primary and secondary antibodies for 3 h using the iBind device. Antibodies included rabbit anti-SARS-CoV-2 S 1:500 (SinoBilogical: 40589-T62), rabbit anti-SARS-CoV-2 N 1:500 (ThermoFisher: TP790189), rabbit anti-SARS-CoV-2 M protein 1:250 (ThermoFisher: PA1-41160), rabbit anti-SARS-CoV-2 E protein 1:250 (ThermoFisher: PA1-41158), rabbit anti-β-actin 1:250 (Abcam: ab197345), and horseradish peroxidase (HRP) conjugated secondary antibody (anti-rabbit IgG) 1:1000. Protein bands were visualised using the Pierce™ ECL Plus Western Blotting Substrate.

### SARS-CoV-2 Transcriptomics

Stable cells expressing SARS-CoV-2 proteins were plated at 4.8×10^5^ cells/well in a 6-well plate and incubated for 48 h. After replacing the media with 500 μl Opti-MEM, cells were rinsed with warm dPBS, and lysed using 350 μl Qiagen buffer RLT with β-mercaptoethanol. RNA purification was performed from cellular lysates using the Qiagen RNAeasy Plus kit, following the manufacturer’s instructions, and subsequently eluted in 35 µl nuclease-free water. Quantification of RNA was carried out using a Qubit high sensitivity RNA fluorometer. PolyA+ mRNA was purified from 35 ng total RNA using the Dynabeads mRNA purification kit and eluted in 15 µl of nuclease-free water. Library preparation involved 30 ng polyA+ mRNA and followed the Oxford Nanopore SQK-PCS-109 protocol, incorporating the EXP-PBC-001 barcoding kit for flow cell multiplexing. Briefly, reverse transcription was conducted using Maxima H Minus Reverse Transcriptase (42 °C for 90 min, 85 °C for 5 min). The resulting reverse transcription product (5 µl) was divided into two 50 µl reactions with Oxford Nanopore barcoded primers for amplification (95 °C for 5 min and 12 cycles of 95 °C for 15 sec, 62 °C for 15 sec, and 65 °C for 8 min, with final elongation at 65 °C for 8 min). Amplification products were pooled and purified using Beckman Coulter Ampure XP magnetic beads with a 0.45x ratio of beads to DNA volume. Library quantity assessment was conducted using Qubit, and each library (150 ng) was sequenced with two samples per flow cell. Sequencing was performed using MIN-106 flow cells (R9.4.1 pores) on the Oxford Nanopore MinION sequencing platform. Run durations ranged from 24-48 h.

### Bioinformatics

Guppy basecaller, configured with a minimum score of 7 and fixed parameters for q-score filtering and barcode kits, was employed to base-call and demultiplex the Fast5 files. An additional parameter was introduced to trim primers and adapters from the sequences. To confirm the expression of the protein of interest, Kraken2 mapped the reads to a viral genome database ^[36]^, while Minimap2 established an index from the Genome Reference Consortium Human Build 38 (hg38). Minimap2 was further utilised to map the reads to the indexed hg38 ^[37]^, and the ’Subread’ function feature of the R package was used for their assignment to genomic features (Liao et al., 2013). Subsequently, EdgeR, transforming raw counts into counts per million (CPM) and applying additional normalisation using trimmed mean M-values (TMM), was employed to normalise raw read counts. The ’voom’ method within Limma was used to fit linear models, identifying differentially expressed genes in the resulting expression file ^[38,39]^. Finally, the normalised count file underwent analysis with the clusterProfiler R package ^[40]^ for gene set enrichment analysis, incorporating Gene Ontology (GO) terms to categorise genes based on biological processes, molecular functions, and cellular components. Additionally, the GSEA software ^[41]^ was employed to assess the enrichment of predefined gene sets. Pathway visualisation utilised the Pathview R package ^[42]^, generating pathway diagrams illustrating affected genes and their interactions within specific biological pathways. Reactome and KEGG databases served as sources of gene sets for the GSEA analysis.

### Statistical Analysis

Statistical analyses were conducted using GraphPad Prism 9.0 software. Unpaired sample comparisons were performed for all data. Non-parametric one-way analysis of variance (ANOVA) using the Kruskal-Wallis test was applied, followed by Dunn’s analysis for paired multiple comparisons. Data are presented as mean ± SEM, with error bars representing the SEM. A significance level of P<0.05 was considered appropriate. Further details about the statistical methods employed can be found in the corresponding figure legend.

## RESULTS

### SARS-CoV-2 Env glycoproteins downmodulate HIV-1 LTR activity

To investigate the impact of SARS-CoV-2 structural proteins (S, M, E, and N) on NF-κB activity we measured luciferase protein expression driven from the HIV-1 subtype B LTR. Co-transfection experiments were conducted using various concentrations of SARS-CoV-2 protein expression plasmids (12, 6, and 1 ng). A fixed amount of reporter LTR plasmid (6 ng) expressing the luciferase protein was used in HEK293T (**Fig 1A**) and BEAS-2B cells (**Fig 1B**). Co-transfections included a Tat protein expression plasmid alongside the LTR reporter plasmid and SARS-CoV-2 expression plasmids to assess their impact on HIV-1 LTR activation measured by luciferase activity (RLUs). Plasmid quantities were normalised using a pCDNA control plasmid in all experiments, and controls included the transfection of the Tat plasmid alone and the LTR reporter plasmid alone.

**Figure 1.**
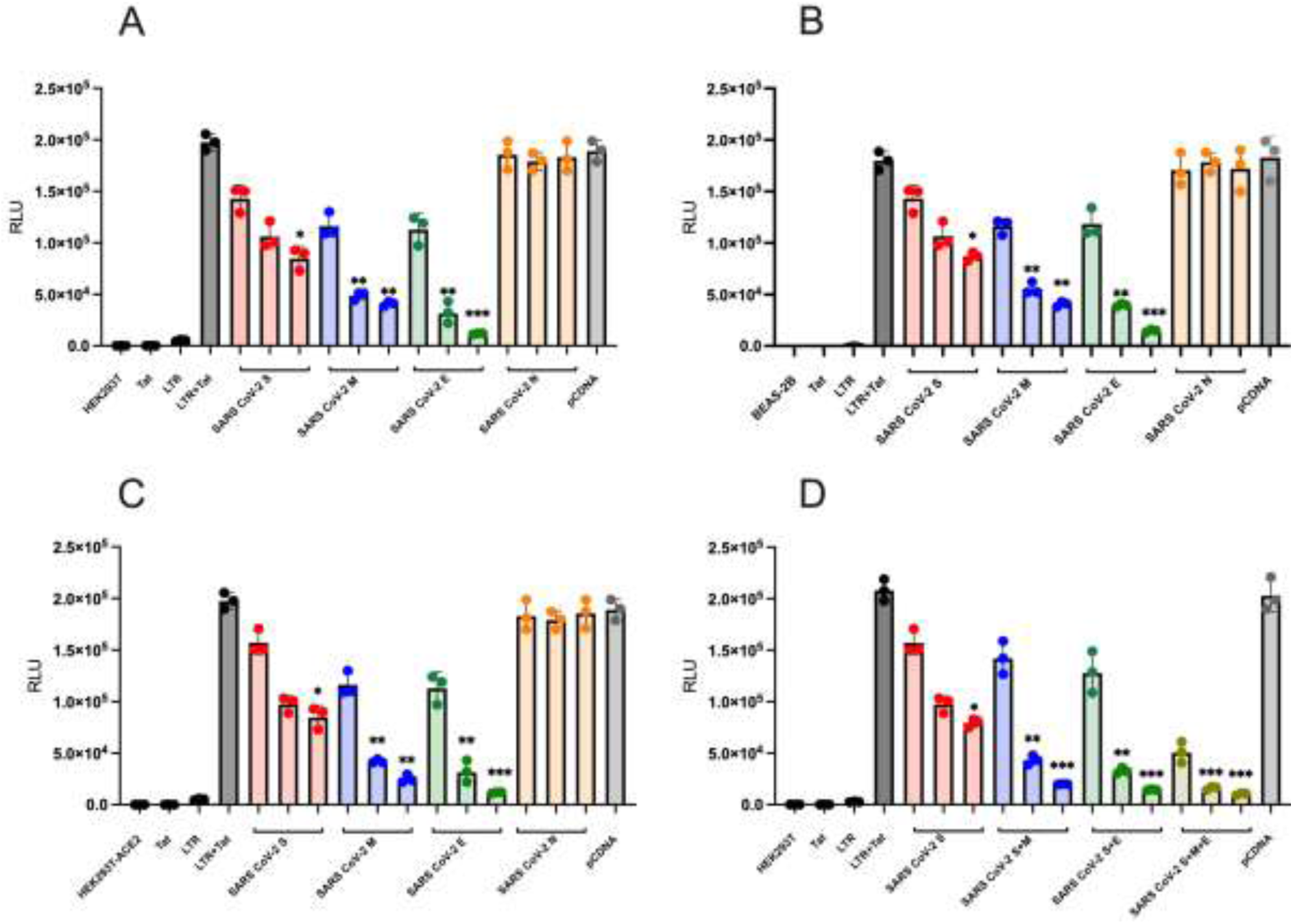
SARS-CoV-2 glycoproteins downmodulate HIV-1 LTR activity. HIV-1 LTR subtype B activation in the presence of varying concentrations of SARS-CoV-2 proteins **(Panel A)** demonstrates the effect of SARS CoV-2 Env glycoproteins on HEK293T cells. **(Panel B)** shows the effect on BEAS-2B cells. **(Panel C)** shows the effect on HEK293T ACE2-expressing cells. **(Panel D)** The effect of combined SARS-CoV-2 S, E and M proteins on HIV-1 LTR activity. Each LTR and Tat was transfected alone as a control for overall LTR activity. Statistical analysis was done using Kruskal-Wallis and Dunn’s tests to compare the control (LTR + Tat) with each concentration of the SARS-CoV-2 proteins and the significance (*P <0.05), (**P <0.01), (***P <0.001) and (ns) represents not significant (p>0.05). The figure illustrates the results of a single experiment, which was conducted in triplicate for validation.

Significant influences of SARS-CoV-2 structural proteins on HIV-1 LTR activity were observed. The S protein demonstrated a dose-dependent effect, inhibitory effect on LTR activation in both cell lines (**Fig.1A and 1B**). At the highest concentration (12 ng) of the S protein, a statistically significant suppression of HIV-1 LTR activity was observed (P<0.05). Similarly, the M protein exhibited a dose-dependent modulation of HIV-1 LTR activity, with statistical significance at concentrations of 12 ng and 6 ng (P<0.01) (**Fig.1A and 1B**). The E protein significantly downregulated HIV-1 LTR activity in both cell lines, particularly at a concentration of 12 ng, with high statistical significance (P<0.001) (**Fig.1A and 1B**). Notably, the inhibitory effect of the E protein surpassed that observed with the S and M proteins. In contrast, the N protein did not affect HIV-1 LTR activity in either cell line tested (**Fig.1A and 1B**).

To explore the potential impact of ACE2 receptor expression on SARS-CoV-2 structural proteins’ ability to suppress HIV-1 LTR activity, co-transfection experiments were conducted using HEK293T-ACE2-expressing cells. These cells were selected as in SARS-CoV-2 infection viral infected cells express the ACE2 receptor which may interact with the structural proteins during synthesis. Results demonstrated that SARS-CoV-2 S, M, and E proteins significantly suppressed HIV-1 LTR activity in HEK293T-ACE2-expressing cells, with no observed effect with the N protein (**Fig.1C**). These effects were consistent with findings in HEK293T cells (**Fig.1A**), suggesting that ACE2 receptor expression does not alter the impact of SARS-CoV-2 S, M, and E proteins on HIV-1 LTR activity.

To investigate the combined effect of SARS-CoV-2 M and E proteins with the S protein on HIV-1 LTR activation, HEK293T were co-transfected with variant concentrations of E and M expression plasmids with or without the S protein. Luciferase activity was quantified to analyse the combined effects of SARS-CoV-2 M, E, and S protein expressions on LTR activation. Combining the SARS-CoV-2 S protein with the M or E proteins had a significant effect on HIV-1 LTR activation, with significant P values (P<0.05) for 12 ng and (P<0.05) for 6 ng (**Fig.1D**). Interestingly, combining all SARS-CoV-2 S, M, and E proteins together had an even greater effect on LTR activation, with both 12 ng and 6 ng showing a significant P value (P<0.05) (**Fig.1D**). These results suggest a synergistic effect of combining multiple viral proteins in enhancing the regulatory function of the S protein on LTR transcriptional activity, highlighting the importance of considering the collective impact of viral proteins in modulating NF-κB activity and potentially HIV-1 infection. These findings provide insights into the interactions between SARS-CoV-2 structural proteins and HIV-1 LTR activity. The S and M proteins exert inhibitory effects, whereas the E protein demonstrates a potent downregulatory effect on LTR activation. The N protein does not appear to impact HIV-1 LTR activity. This study contributes to our understanding of the molecular interplay between SARS-CoV-2 and HIV-1, offering insights into potential mechanisms underlying the modulation of HIV-1 replication by SARS-CoV-2 structural proteins.

### Impact of S Protein Variants on Down-modulation of HIV-1 LTR Activity and Mutation Influence

Our initial analysis revealed a concentration-dependent down-modulation of HIV-1 LTR activity by the SARS-CoV-2 S protein, as evidenced by a reduction in luciferase activity at higher concentrations. To delve deeper into the influence of S protein variants on their ability to downmodulate HIV-1 LTR activity, we conducted experiments in HEK293T cells. These cells were transfected with varying concentrations of SARS-CoV-2 plasmids expressing S Env proteins representing different viral strains (Alpha, Beta, Gamma, Delta, and Omicron), along with LTR luciferase reporter and Tat expression plasmids. Equal plasmid quantities were maintained across all experiments, with pcDNA serving to standardise DNA transfection levels. To ensure specificity in the interaction between the S protein and HIV-1 LTR, control experiments included transfections with only the Tat expression plasmid or the HIV-1 LTR reporter plasmid. Luciferase activity was quantified by measuring relative light units (RLUs) in all conditions. Our findings indicate that the Alpha variant exerts a comparable effect on downmodulating HIV-1 LTR activity as the SARS-CoV-2 Wuhan S protein, with significant down-modulation observed at a concentration of 12 ng (P<0.05). Conversely, the Beta, Gamma, and Delta variants inhibited HIV-1 LTR activity, although the observed down-modulation did not reach statistical significance when compared to the Alpha variant **(Fig.2A)**. Intriguingly, the Omicron variant demonstrated no effect on HIV-1 LTR activity. Notably, the Omicron variant harbours a significantly larger number of mutations in the S protein than the other variants **(Fig.2B)**. This implies that SARS-CoV-2 S variants have evolved from suppressing NF-κB signalling and potentially altering immune induction.

**Figure 2.**
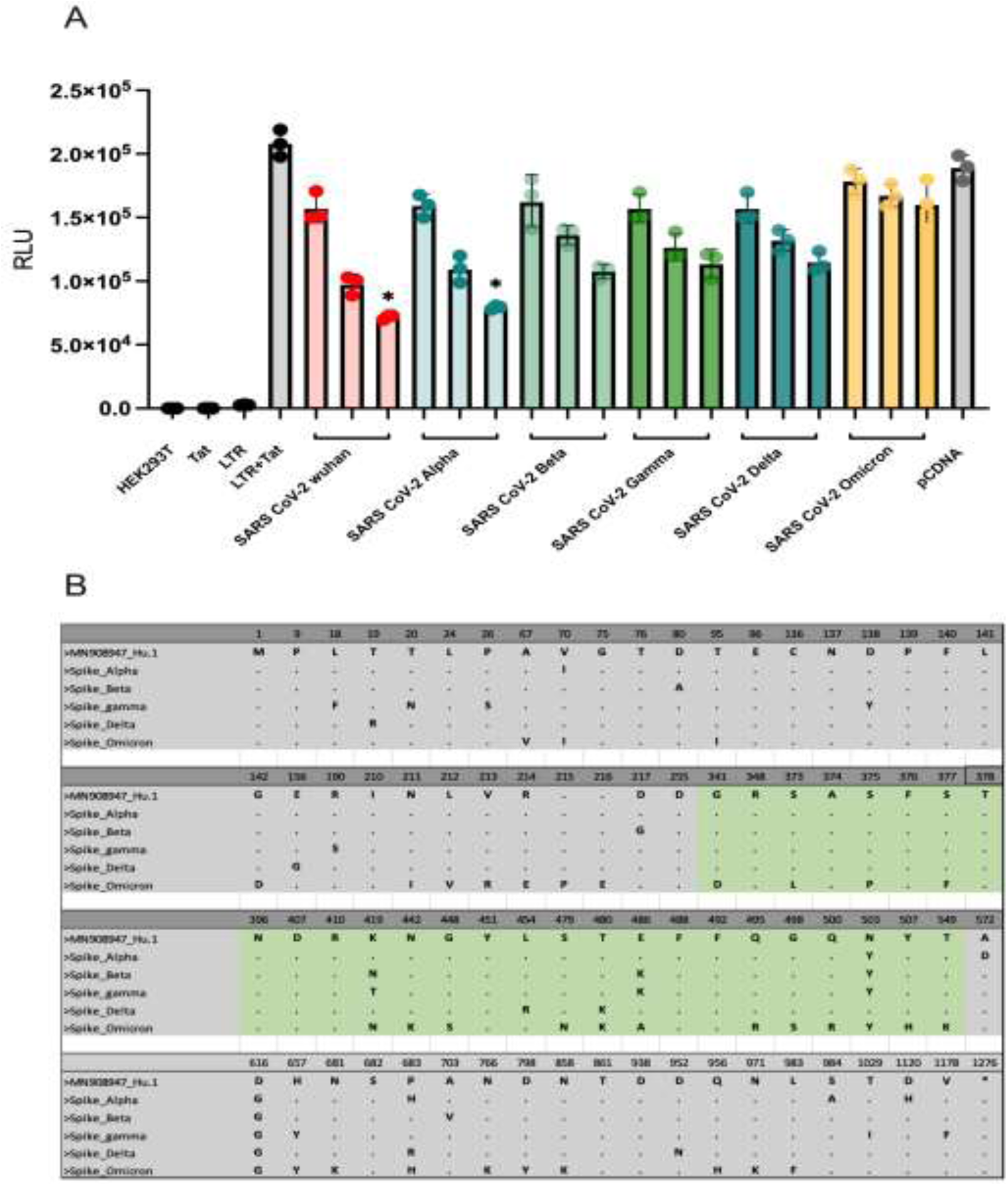
The impact of spike protein mutations on its ability to downmodulate HIV-1 LTR activity. **(Panel A)** depicts the activation of HIV-1 LTR subtype B when co-transfected with varying concentrations of the SARS-CoV-2 wuhan spike protein, Alpha, Beta, Gamma, Delta and Omicron variants, Each LTR and Tat was transfected alone as a control for overall LTR activity. Kruskal-Wallis and Dunn’s tests were used to perform a statistical comparison between the control (LTR + Tat) and each concentration of SARS-CoV-2 spike variants protein. The P value for the concentration of 12ng of SARS-CoV-2 spike and Alpha variants were found to be (*P <0.05). **(Panel B)** Comparative sequence alignment of spike protein amino acid sequences among SARS-CoV-2 wild type and Alpha, Beta, Delta, Gamma, and Omicron variants. The alignment highlights the positions where variation has occurred in the spike protein when compared to the sequence from the strain (MN908947_Hu.1) first isolated in humans in Wuhan/China. The numbering above indicates the amino-acid position within the full-length spike protein spanning from 1 to 1275. The green highlights indicate the variant amino-acid positions within the receptor binding domain (RBD) region.

### SARS-CoV-2 Differential Gene Expression Analysis

The introduction of SARS-CoV-2 structural proteins (S, M, and E) led to the down-regulation of HIV-1 LTR activation. The study aimed to elucidate the mRNA transcriptional profile of cells expressing these proteins, hypothesising an impact on cellular transcription factors like NF-κB. To achieve this, a comprehensive differential gene expression (DGE) analysis was conducted, utilising HEK293T cells stably transduced with SARS-CoV-2 S, M, E, and N proteins, with mock cells as a control group. Three replicates were employed for each condition (Mock n= 3, CoV-2 S n= 3, CoV-2 M n= 3, CoV-2 E n= 3, and CoV-2 N n= 3). Total RNA extraction, reverse transcription, and generation of barcoded cDNA libraries through PCR amplification were conducted. Subsequently, the libraries were sequenced on the ONT’s MinION platform, with two libraries per flow cell. R packages, including ’EdgeR’ and ’Limma,’ were employed for analysis, encompassing barcode and adapter trimming. The sequenced libraries demonstrated consistent characteristics across read count, average read length, and N50 value (data not shown). Verification of SARS-CoV-2 protein presence in HEK293T cells utilised Kraken2, revealing varying percentages of reads classified as SARS-CoV-2: 0.5% for S and N samples, 2.8% for M samples, and 1% for E samples, while mock samples showed no SARS-CoV-2 reads (data not shown). Alignment of libraries to the human genome (hg38) and assignment of genomic features facilitated raw gene count generation. DGE analysis involved EdgeR transformation, normalisation using trimmed mean M-values (TMM), and subsequent analysis using Limma. Genes with low expression levels, which may not provide meaningful biological insights, were filtered out using the EdgeR function ’filterByExpr’ with default parameters. This filtering step resulted in a library that facilitated the comparison of gene expression, as depicted in (data not shown).

The libraries were normalised using the EdgeR function, applying the trimmed mean M-values (TMM) method. This method determined scaling factors based on the sample with a library size closest to the mean of all samples (data not shown). The normalised and filtered expression library underwent DGE analysis using the Limma package in R. The Voom function in Limma was initially employed to transform the dataset from counts per million (CPM) to log2 CPM, generating a curve representing the relationship between the square root of the standard deviation of mean log2 CPM and mean expression. This curve demonstrated a decrease in genewise variance with increasing mean expression (data not shown). Subsequently, Limma was utilised to compare gene expression between mock and SARS-CoV-2 samples using empirical Bayes models. These models effectively reduced the residual variance of genes with high and low variance, enhancing statistical power by shrinking them towards the average (data not shown). This adjustment ensured that genewise variance was no longer dependent on mean expression levels, providing a robust foundation for the subsequent analyses. The Limma analysis outcomes, including normalised expression levels, log2 fold change (logFC), and p-values, were further investigated. Volcano plots illustrated differential gene expression between mock and SARS-CoV-2 S, M, E, and N transduced cells. Significance criteria were set at a log10 adjusted p-value of 1.3 (p=0.05) and a logFC of 1 or -1 (equivalent to a 2-fold difference between conditions). Genes meeting both criteria were deemed significantly upregulated or downregulated in the presence of SARS-CoV-2 proteins (**Fig.3A-3D**). According to the volcano plots, numerous genes related to interferon-stimulated genes (ISGs), 2′-5′-oligoadenylate synthetases (OASs), heat shock proteins (HSPs), co-chaperones, and certain transcription factors were significantly upregulated in the presence of SARS-CoV-2 proteins. Conversely, long non-coding RNAs (lncRNAs), transcription factors, and immune response-related genes were notably downregulated in the presence of SARS-CoV-2 proteins, with specific patterns observed for different structural proteins.

**Figure 3.**
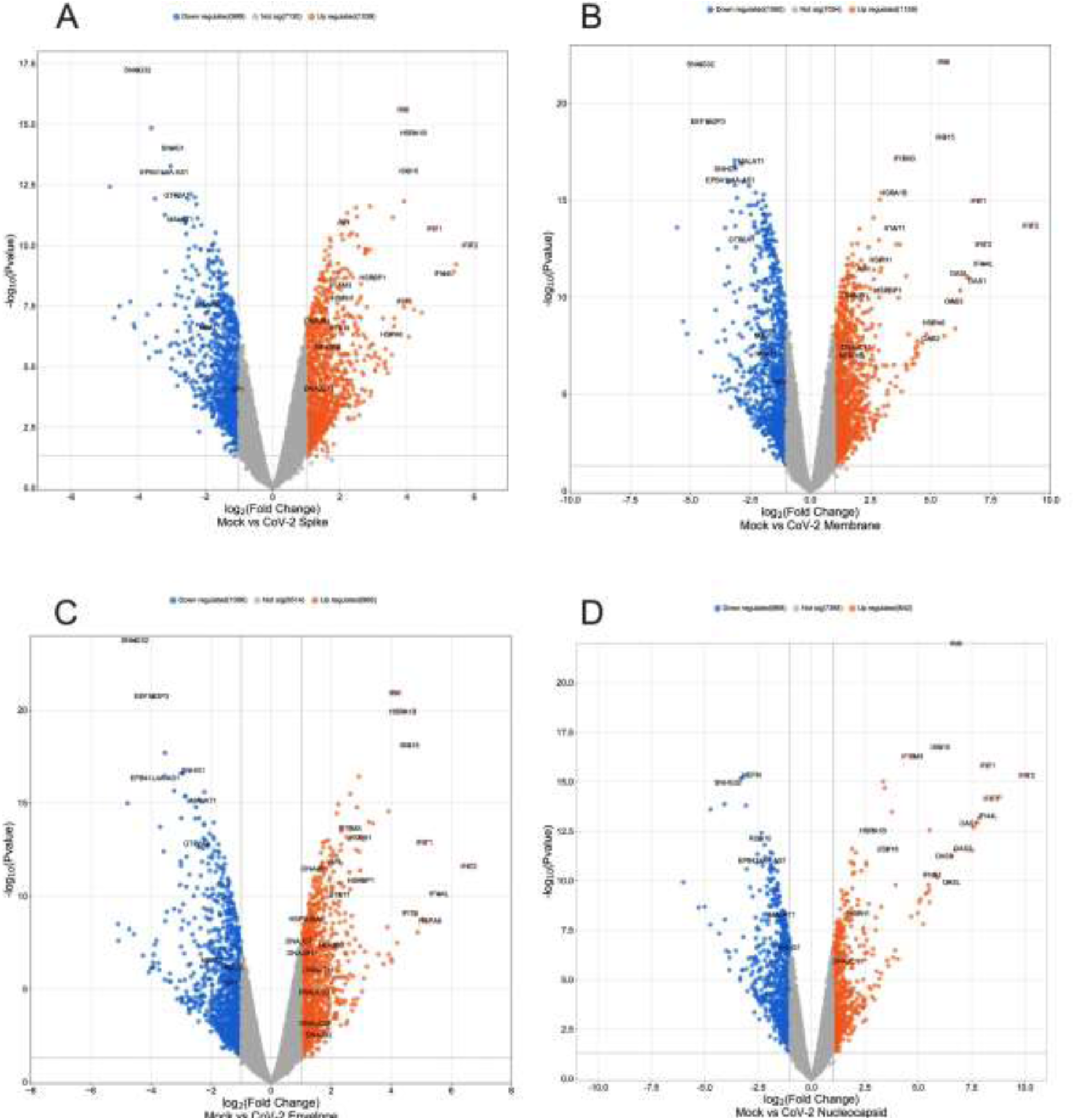
Volcano plots of differential gene expression analysis in SARS CoV-2 transcriptomics. Each panel **(A-D)** corresponds to a specific viral structural protein: **(A)** Spike, **(B)** Membrane, **(C)** Envelope, and **(D)** Nucleocapsid. Volcano Plot displaying -Log2 adjusted P value versus Log2 fold change for each gene. Dashed lines indicate the thresholds, with a P value greater than 1.3 -Log10 and a fold change greater than 1 or less than -1. The gene names of the most substantially differentially expressed genes are annotated.

### SARS-CoV-2 proteins induced interferon-stimulated genes (ISGs)

To assess the expression of key genes associated with ISGs across various groups of transduced cells, normalised counts for each gene, expressed as counts per million (CPM), were graphically represented. This analytical approach facilitated a quantitative evaluation of gene expression levels under different experimental conditions. The ensuing results demonstrated a significant increase in the expression of several ISGs, including ISG15, IFI6, IFIT1, IFIT2, IFIT3, IFI44L, and IFITM3, in the presence of SARS-CoV-2 S, M, E, and N proteins, as illustrated in (**Fig.4**). These expression profiles underscore the virus’s capacity to upregulate ISGs. The observed elevation of ISGs, encompassing ISG15, IFI6, IFIT1, IFIT2, IFIT3, IFI44L, and IFITM3, highlights SARS-CoV-2’s ability to activate the host’s interferon response—a critical defence mechanism against viral infection.

**Figure 4.**
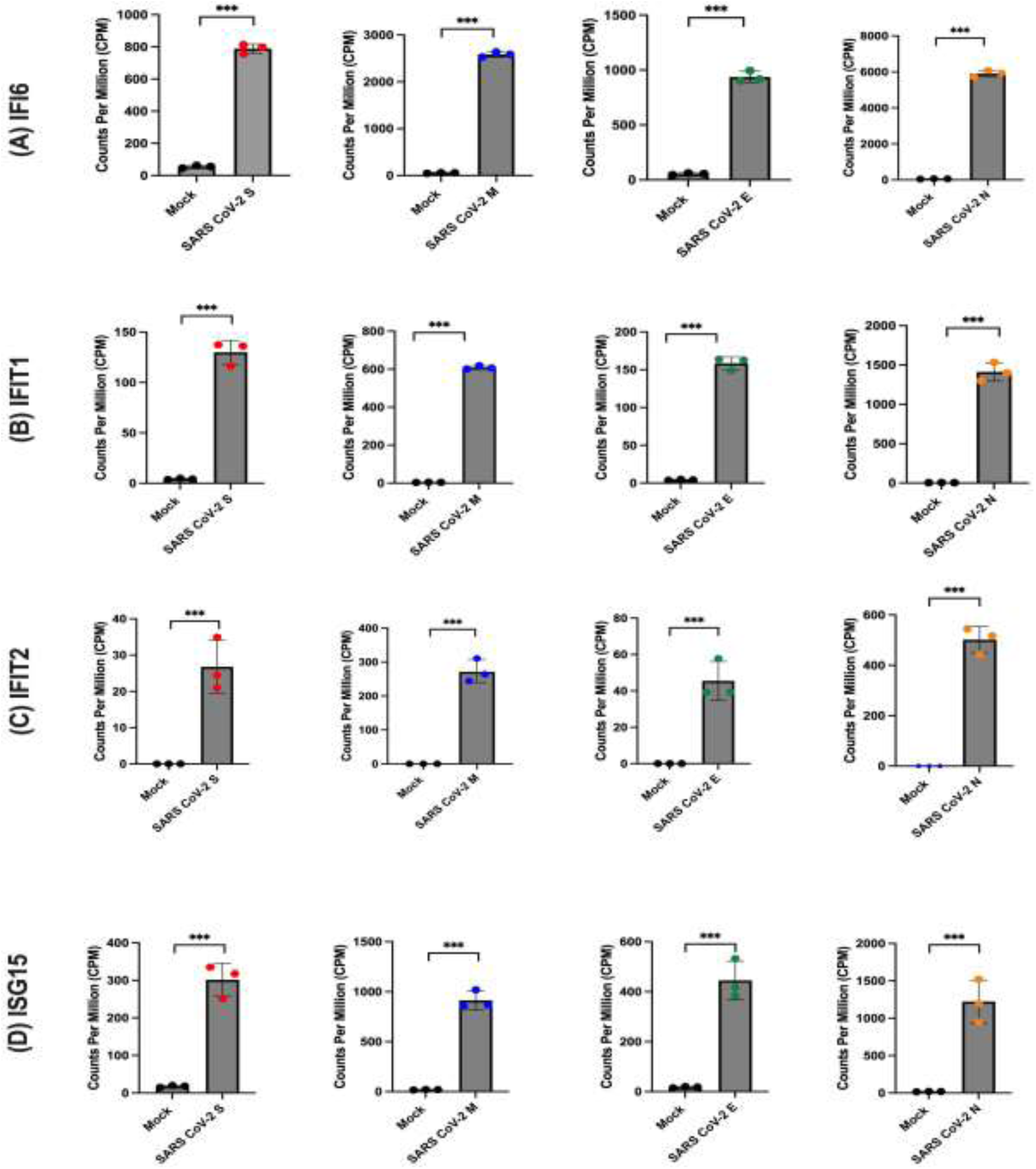
Expression levels of genes associated with interferon-stimulated genes (ISGs) in the presence or absence of SARS CoV-2 proteins. The normalised counts, expressed as counts per million (CPM), are plotted for each sample group, including Mock (n=3), SARS-CoV-2 S (n=3), SARS-CoV-2 M (n=3), SARS-CoV-2 E (n=3), and SARS-CoV-2 N (n=3). Panel (**A**) presents the comparison of CPM for IFI16, panel (**B**) for IFIT1, panel (**C**) for IFIT2, panel (**D**) for ISG15. The statistical significance of differential gene expression was determined using the Voom/Limma analysis, and the results were indicated as ns (not significant), *P < 0.05, **P < 0.01, or ***P < 0.001.

### SARS-CoV-2 proteins induced heat shock proteins (HSPs) and co-chaperones

The analysis of differential gene expression (DGE) revealed that SARS-CoV-2 structural proteins induce significant up-regulation of heat shock proteins (HSPs) and co-chaperones. Notably, HSPA6, HSPA1B, HSPBP1, and HSPH1 were identified as HSPs with increased expression. However, the SARS-CoV-2 N protein showed comparatively less stimulation of HSPs compared to other structural proteins. In comparison to mock conditions, the presence of SARS-CoV-2 N protein significantly upregulated only HSPA1B and HSPH1, while HSPBP1 exhibited weaker up-regulation with the N protein than other structural proteins (**Fig 5**). These findings provide insights into how SARS-CoV-2 interacts with cellular stress response mechanisms through HSPs, suggesting their involvement in cellular stress pathways. The differential stimulation of HSPs by specific viral proteins implies distinct mechanisms by which different SARS-CoV-2 components interact with host cellular machinery.

**Figure 5.**
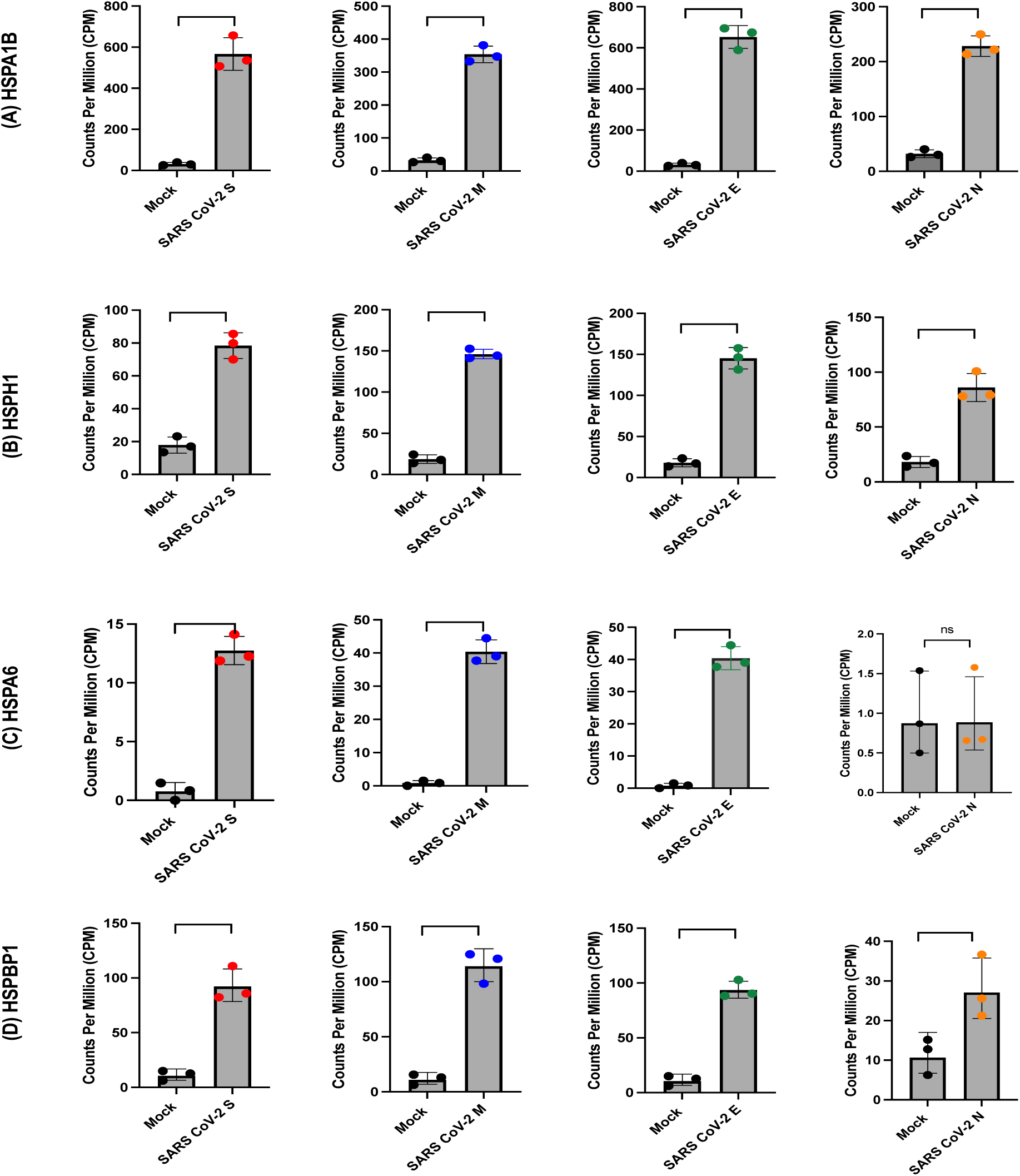
Expression levels of genes associated with heat shock proteins (HSPs) in the presence or absence of SARS-CoV-2 proteins. The normalised counts, expressed as counts per million (CPM), are plotted for each sample group, including Mock (n=3), SARS-CoV-2 S (n=3), SARS-CoV-2 M (n=3), SARS-CoV-2 E (n=3), and SARS-CoV-2 N (n=3). Panel (**A**) displays the comparison of CPM for HSPA1B, panel (**B**) for HSPH1, panel (**C**) for HSPA6, panel (**D**) for HSPBP1. The statistical significance of differential gene expression was determined using the Voom/Limma analysis, and the results were indicated as ns (not significant), *P < 0.05, **P < 0.01, or ***P < 0.001.

### Expression of SARS-CoV-2 proteins regulates long non-coding RNAs (lncRNAs)

The assessment of gene expression profiles related to long non-coding RNAs (lncRNAs) in cells expressing SARS-CoV-2 proteins involved analysing counts per million (CPM) values for the most significantly different genes. As illustrated in (**Fig.6**), the results indicated a significant decrease in the expression of SNHG1, SNHG32, EPB41L4A-AS1, and MALAT1 in the presence of SARS-CoV-2 proteins. This observation suggests that SARS-CoV-2 proteins negatively influence the expression of these lncRNA-associated genes, revealing a potential regulatory mechanism employed by the virus to modulate host cellular processes through intricate interactions with the transcriptional machinery.

**Figure 6.**
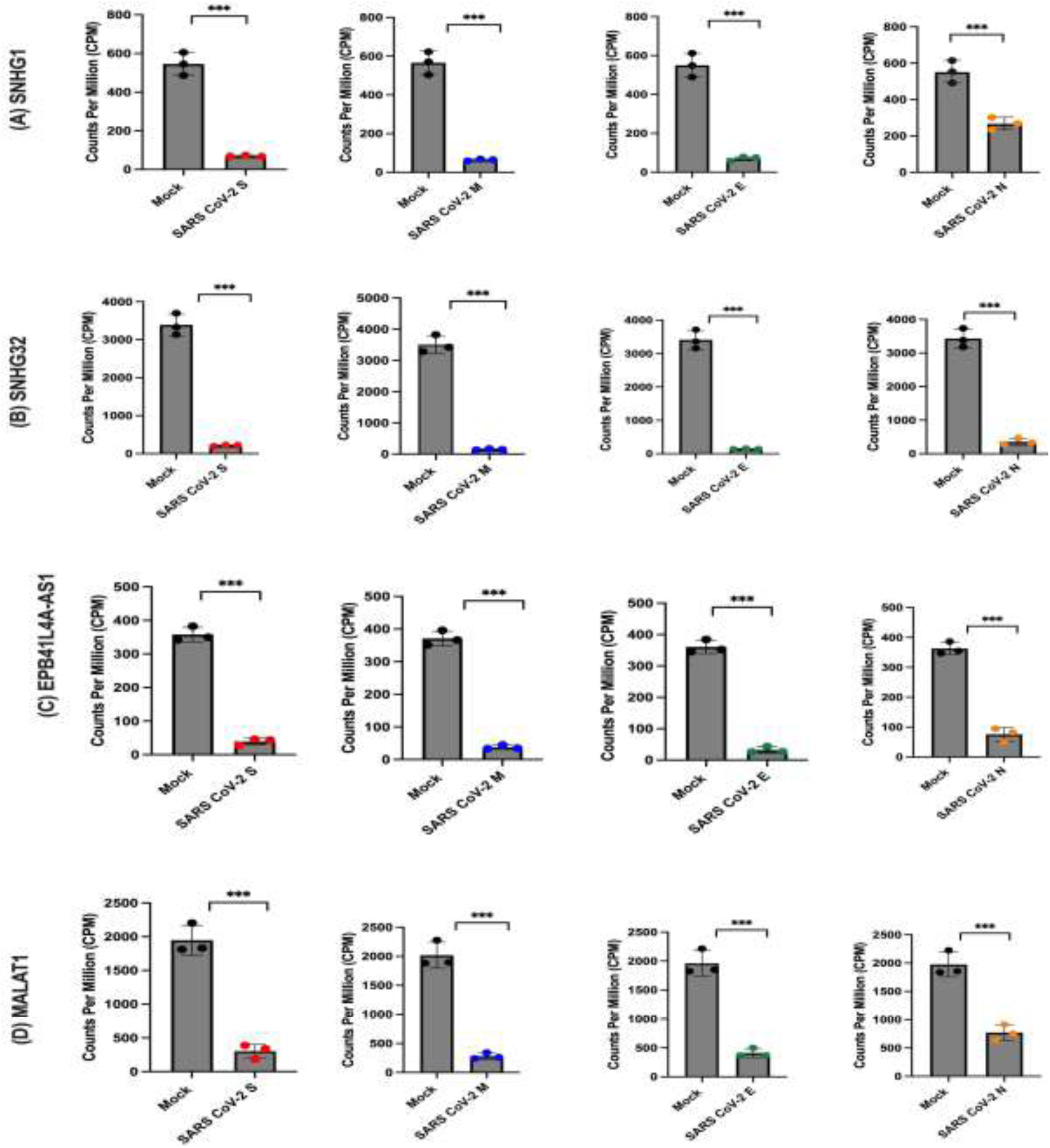
Expression levels of genes associated with long non-coding RNAs (lncRNAs) in the presence or absence of SARS-CoV-2 structural proteins. The normalised counts, expressed as counts per million (CPM), are plotted for each sample group, including Mock (n=3), SARS-CoV-2 S (n=3), SARS-CoV-2 M (n=3), SARS-CoV-2 E (n=3), and SARS-CoV-2 N (n=3). Panel (**A**) displays the comparison of CPM for SNHG1, panel (**B**) for SNHG32, panel (**C**) for EPB4IL4A-AS1, and panel (**D**) for MALAT1. The statistical significance of differential gene expression was determined using the Voom/Limma analysis, and the results were indicated as ns (not significant), *P < 0.05, **P < 0.01, or ***P < 0.001.

### SARS CoV-2 proteins targeting transcription factors

In comparing the transcriptome of cells expressing SARS-CoV-2 structural proteins to a mock condition, our DGE analysis revealed alterations in various host transcription factors. We focused on the expression levels of NF-κB1, SP1, JUN, STAT1, and GTF2A1, which were among the most significantly differentially expressed factors (**Fig.7**). Notably, NF-κB1 did not show a significant difference in expression in the presence of SARS-CoV-2 proteins, but we observed an up-regulation of its inhibitor, NFκBIB (Figure 5.14A). Furthermore, SP1 and GTF2A1 displayed substantial down-regulation in the presence of SARS-CoV-2 S, M, and E proteins, indicating a potential direct influence of these viral proteins on their expression and suggesting an impact on transcription regulation. Conversely, JUN exhibited significant up-regulation, implying its involvement in the cellular response to SARS-CoV-2 infection. Additionally, STAT1 showed significant up-regulation with all SARS-CoV-2 structural proteins compared to the mock condition, suggesting a role in the host antiviral response by potentially mediating downstream signalling pathways triggered by SARS-CoV-2 infection. Our findings indicate that SARS-CoV-2 proteins have the capacity to modulate the expression of specific transcription factors, potentially influencing host-virus interactions and the host’s antiviral response.

**Figure 7.**
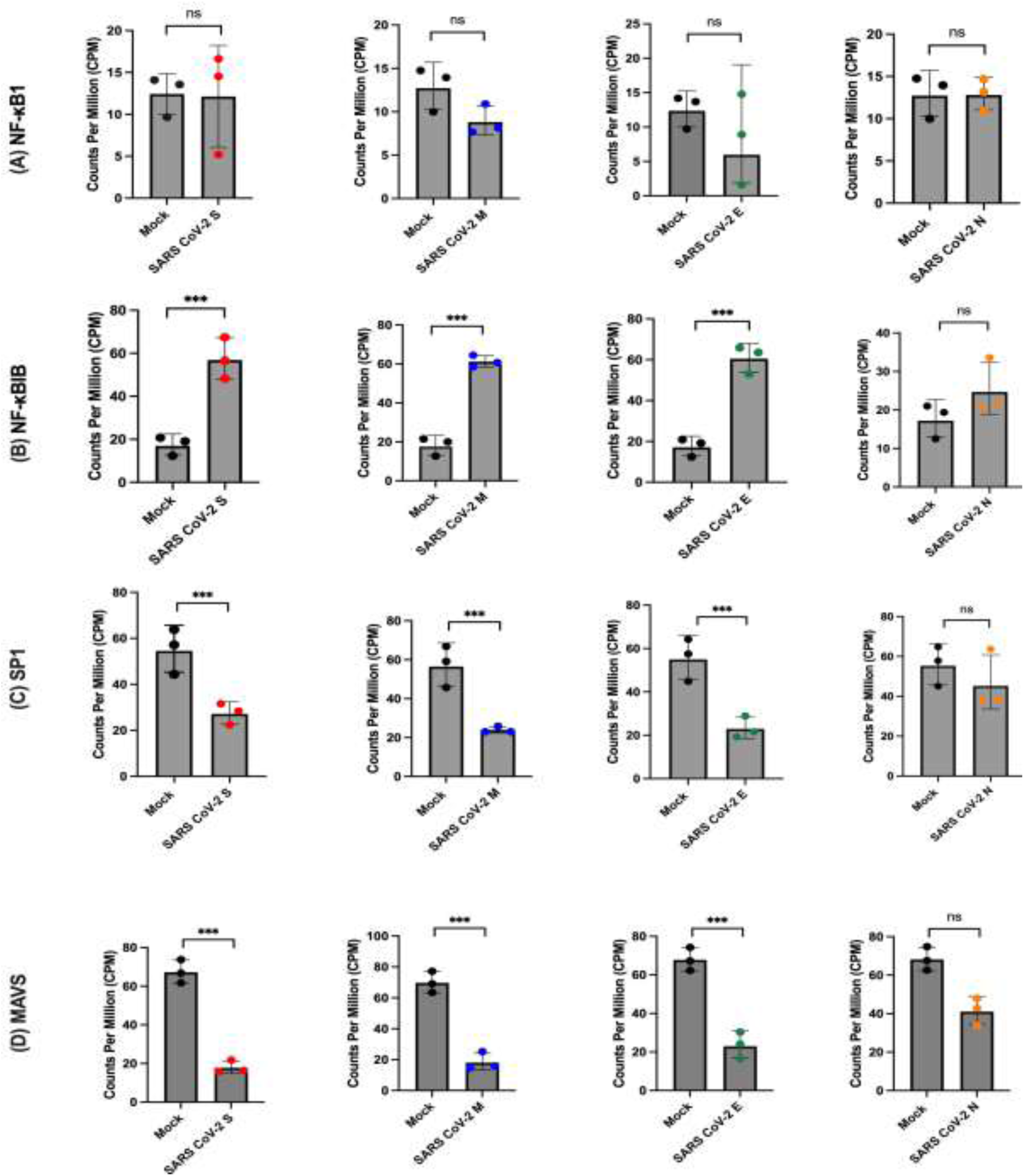
Expression of viral transcription factors in the presence or absence of SARS CoV-2 proteins. Normalised counts for each gene, expressed as counts per million (CPM), are plotted for each sample group (Mock n= 4, SARS CoV-2 n= 3). Panel (**A**) displays the comparison of CPM for for NF-κB1. (**B**) for NF-κBIB (**C**) for SP1 (**D**) For MAVS. The statistical significance of differential gene expression was determined using the Voom/Limma analysis, and the results were indicated as ns (not significant), *P < 0.05, **P < 0.01, or ***P < 0.001.

### Pathway enrichment analysis of SARS CoV-2 transcriptome datasets

We conducted Gene Ontology (GO) and Gene Set Enrichment Analysis (GSEA) on up/downregulated DGE datasets for SARS-CoV-2 S, M, E, and N transcriptomes compared to control. The analysis encompassed the top 10 enriched terms in biological processes (BP), cellular components (CC), and molecular functions (MF). Focusing on biological processes (BP), upregulated DGEs in SARS-CoV-2 S, M, and E datasets exhibited significant enrichment in processes such as protein localisation to the endoplasmic reticulum (ER) and membrane, co-translational protein targeting, translocation of proteins during translation initiation to the ER, cellular response to interferon type I, defence response to viruses, viral gene expression, and nuclear-transcribed mRNA catabolic process (**Fig.8A-C**). Conversely, the SARS-CoV-2 N dataset (**Fig.8D**) displayed notable differences, as ER-related processes were not significantly enriched. Enriched biological processes in this dataset included the response to interferon type I, defence response to viruses, viral gene expression, rRNA processing, and nuclear-transcribed mRNA catabolic process. Additionally, the downregulated DGEs (**Fig.9A-D**) shared common BP terms enriched in viral transcription, translational initiation, non-coding RNA metabolism, ribosome biogenesis, rRNA processing, and ion transport regulation across S, M, E, and N datasets.

**Figure 8.**
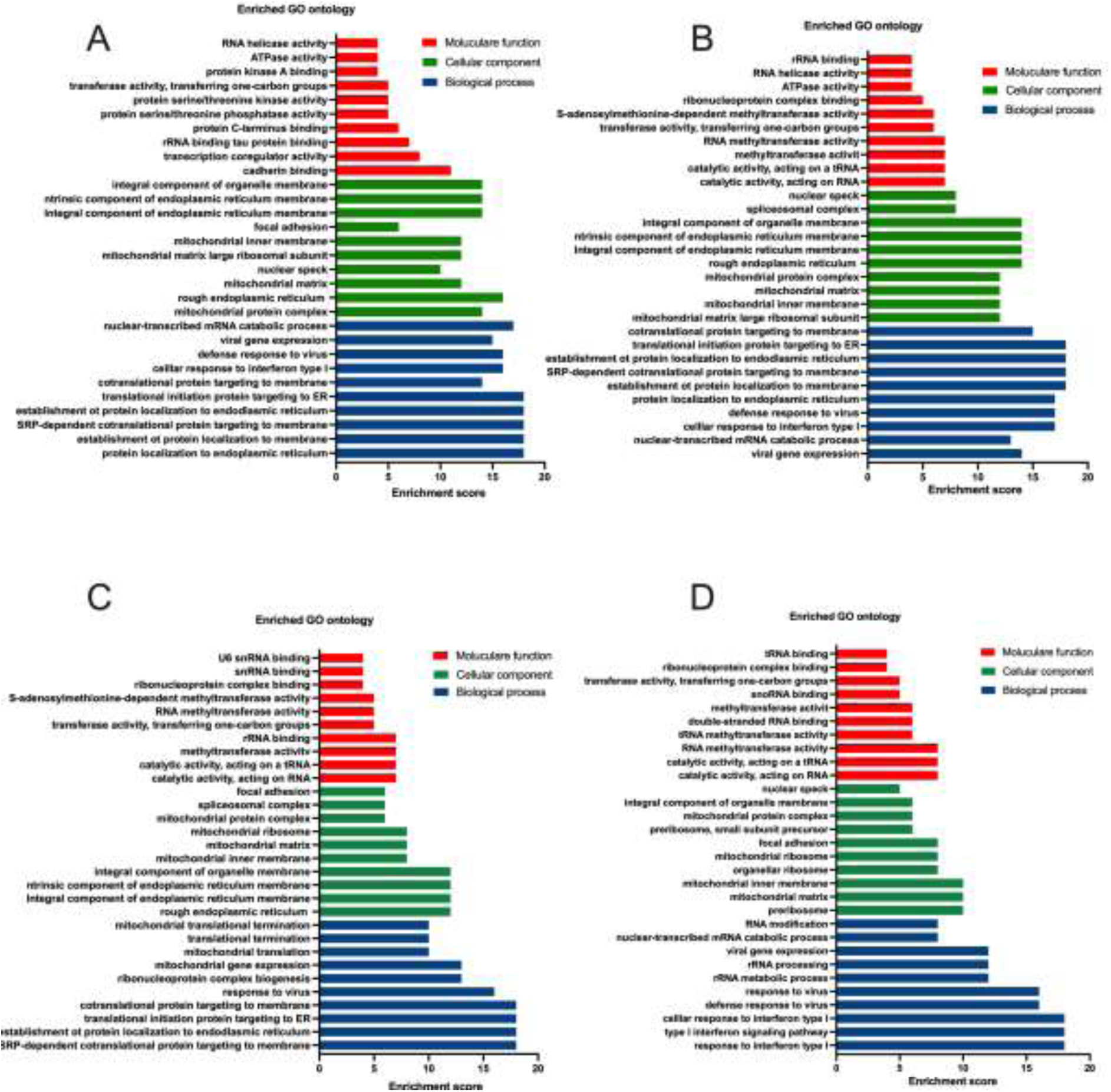
Gene ontology (GO) enrichment analysis for up-regulation DGEs in SARS CoV-2 spike, membrane, envelope and nucleocapsid transcriptome. GO annotations showing top 10 terms significant enrichment of three main categories (biological process, cellular component, and molecular function) with the adjusted p-value <0.05. (**A**) SARS CoV-2 S, (**B**), SARS CoV-2 M, (**C**) SARS CoV-2 E, (**D**) SARS CoV-2 N.

**Figure 9.**
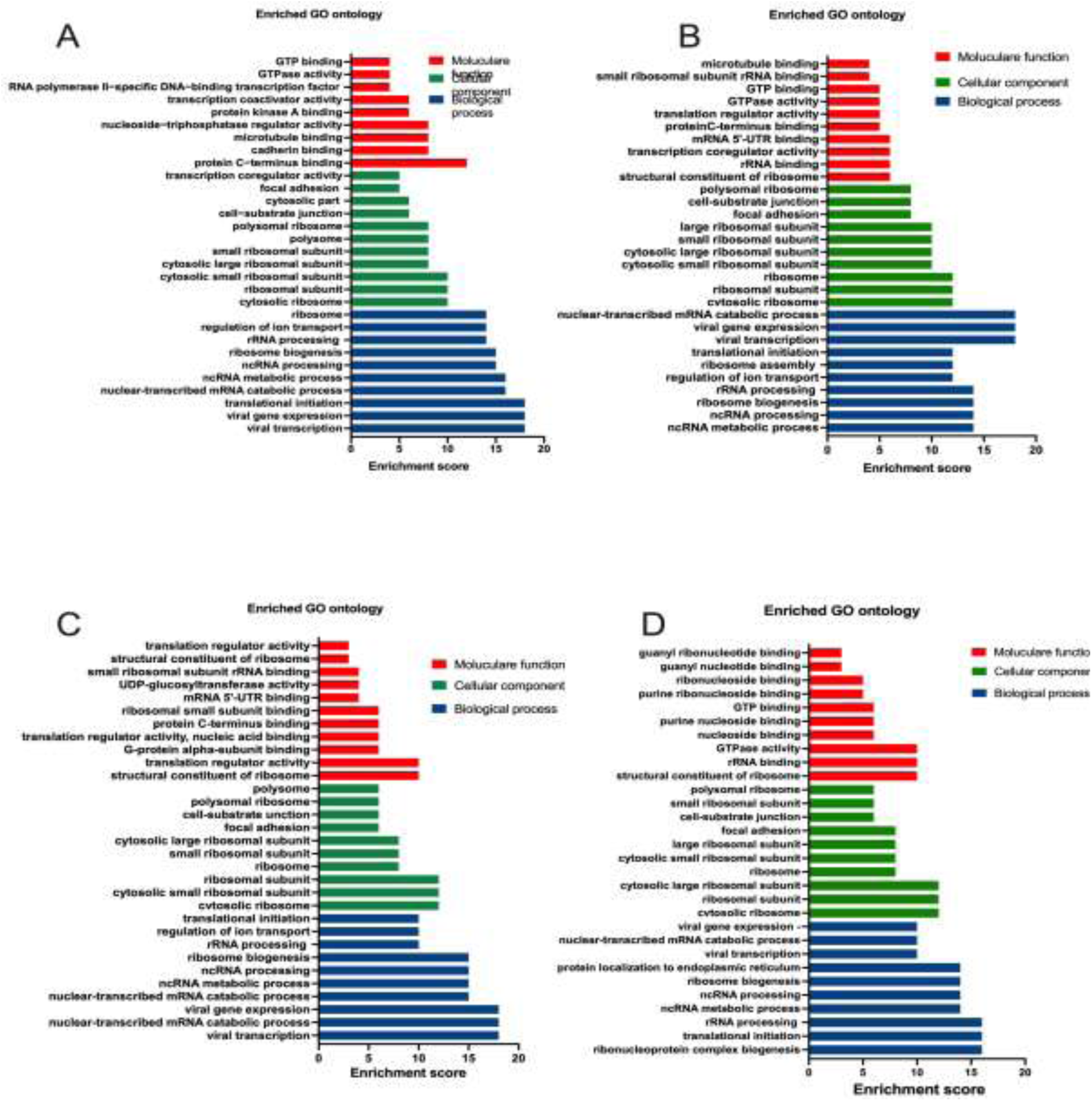
Gene ontology (GO) enrichment analysis for down-regulation DGEs in SARS CoV-2 spike, membrane, envelope and nucleocapsid transcriptome. Top 10 terms were plotted for biological Process, molecular function and cellular component terms *in* SARS CoV-2 S (A), SARS CoV-2 M (B), SARS CoV-2 E (C), and SARS CoV-2 N (D).

## DISCUSSION

Despite more than three years since the onset of the COVID-19 pandemic, several crucial questions regarding SARS-CoV-2 infection and the wide range of symptoms it induces remain unanswered. Among these pertains to the interplay between SARS-CoV-2 and the host during infection. The findings here demonstrate that SARS CoV-2 S, M and E Env proteins can down-modulate HIV-1 LTR activity, similar to the observed effects of the HCV E1E2 and sE2 Env proteins ^[34]^. Both SARS CoV-2 and HCV are positive-sense RNA viruses that replicate in the cytoplasm and have a similar mechanism of viral assembly and release. The fact that SARS CoV-2 S, M, and E proteins have the same capacity to modulate HIV-1 LTR suggests that this effect could be a result of stimulating the ER stress pathway, which was previously observed in the context of HCV infection. It is well known that the SARS CoV-2 Env proteins are synthesized, inserted, and folded in the ER ^[43,44,9]^ and many studies have suggested that it can induce ER stress responses ^[45,46]^. The activation of ER stress can result in the up-regulation of genes involved in protein folding, degradation, and secretion, which may explain the observed down-modulation of HIV-1 LTR activity induced by SARS CoV-2 S, M, and E protein expression. The ability of SARS-CoV-2 S, M and E Env proteins to downmodulate HIV-1 LTR activity may suggest that these proteins play a role in the regulation of host gene expression, potentially contributing to immune evasion and/or pathogenicity. In contrast, the SARS CoV-2 N Env protein is synthesized in the cytoplasm of infected cells ^[47,48]^, unlike the other SARS CoV-2 structural proteins (S, M and E). The finding that the SARS-CoV-2 N protein had no effect on modulating HIV-1 LTR activity may be due to its synthesis and assembly outside the ER, limiting its ability to trigger ER stress.

Here a separate and comparative transcriptome analysis was conducted on cells stably expressing individual SARS-CoV-2 structural proteins (S, M, E and N) compared to control cells to unravel the intricate molecular interactions between viral proteins and host cell machinery, as well as to identify the pathways affected by these protein-induced changes. Interferons (IFNs) are crucial for the induction of host immune responses against viruses. They alert cells to a viral infection and initiate an antiviral response. Infected cells release type I interferons (IFN-alpha and IFN-beta), which induce the expression of interferon-stimulated genes (ISGs) ^[49]^. ISGs produce proteins that inhibit viral replication, activate immune cells, and enhance overall host defence mechanisms ^[50,49]^. In the context of SARS-CoV-2 infection, a robust immune response is triggered, leading to the induction of type I interferons (IFNβ and IFNα) and type III interferon (IFNλ) in various cell types ^[51–54]^. Furthermore, RNA sequencing analysis of nasopharyngeal swabs from COVID-19 patients with diverse disease profiles has confirmed the up-regulation of key interferon-stimulated genes (ISGs) ^[55]^. However, despite the induction of interferon responses by related coronaviruses such as SARS-CoV and MERS-CoV, emerging evidence suggests that SARS-CoV-2 may have evolved mechanisms to dampen the production and signalling of type I interferons, allowing the virus to replicate and spread within the host without eliciting an effective antiviral response ^[56,57]^. In a recent study, 23 viral proteins were screened, revealing that SARS-CoV-2 NSP1, NSP3, NSP12, NSP13, NSP14, ORF3, ORF6, and M protein inhibit Sendai virus-induced IFN-β promoter activation, while NSP2 and S protein have the opposite effect ^[58]^.

The analysis of DGEs demonstrated that expressing SARS-CoV-2 structural proteins induces the up-regulation of specific interferon-stimulated genes (ISGs) such as ISG15, IFI6, IFIT1, IFIT2, IFIT3, IFI44L, IFITM3, and members of the OAS family. These findings indicate that the SARS-CoV-2 proteins have diverse functions in modulating the host’s innate immune response. Similar, a transcriptomic analysis conducted on SARS-CoV-2-infected calu3 cells demonstrated an up-regulation of genes associated with the innate immune response, including IFIT2, OAS2, and IFNB1 ^[59]^. Over the course of the COVID-19 pandemic, numerous studies have investigated the inflammatory response induced by coronavirus infection, shedding light on novel roles of heat shock proteins (HSPs) in this context ^[60,61]^. Increased expression of HSPs may play a role in protecting cells from stress induced by the viral infection and aiding in the proper folding and stability of proteins ^[62]^. In our study, among the significant upregulated DGEs, many are HSPs encoding genes such as HSPA6, HSPA1B, HSPBP1 HSPH1. However, the stimulation of HSPs by the SARS-CoV-2 N protein was comparatively lower than that observed with other SARS-CoV-2 structural proteins. In addition, DNAJB1 and DNAJC11 were significant up-regulated in response to SARS-CoV-2 structural proteins. In a similar vein, a study investigating the transcriptome of SARS-CoV-2-infected cells revealed a substantial up-regulation of HSP-encoding genes, including HSPA6 and HSPA1B ^[63]^. Furthermore, the expression of DNAJB1 and DNAJB9, members of the DnaJ homolog subfamily B, was also found to be elevated in response to SARS-CoV-2 infection ^[63]^. These findings support our current study, highlighting the consistent involvement of HSPs and molecular chaperones in the cellular response to SARS-CoV-2 proteins. Further, we observed significant down-regulation of lncRNAs related genes including SNHG1 and SNHG32, EPB41L4A-AS1, MALAT1 and NEAT1 in the precent of SARS CoV-2 structural proteins. As described above, lncRNAs are emerging as regulatory factors involved in diverse biological processes and have been implicated in the innate immune response, potentially through their interactions with IFN-related pathways ^[64–66]^. Emerging evidence suggests that certain long non-coding RNAs (lncRNAs), such as metastasis-associated lung adenocarcinoma transcript 1 (MALAT1) and nuclear paraspeckle assembly transcript 1 (NEAT1), are closely linked to immune responses and may contribute to the inflammatory processes observed in SARS-CoV-2 infected cells ^[67,68]^. Notably, viral infection-induced down-regulation of MALAT1 expression has been associated with enhanced activation of IRF3 and production of type I IFNs, suggesting its antiviral role in therapy ^[69]^. However, our current understanding of the specific functions of lncRNAs in innate immune regulation related to interferons in COVID-19 patients is limited. Additionally, we observed alterations in the expression of various transcription factors upon exposure to SARS-CoV-2 structural proteins. Specifically, we observed up-regulation of JUN and STAT1, while NF-κB1 remained unchanged. Interestingly, the inhibitor of NF-κB, NFκBIB were shown to be upregulated. Additionally, we observed down-regulation of SP1 and GTF2A1. Notably, the influence of the N protein was observed only on the up-regulation of STAT1. These findings suggest that SARS-CoV-2 structural proteins have selective effects on the expression of transcription factors, potentially impacting the transcriptional regulation of target genes involved in immune responses and other biological processes. It is worth mentioning that in the presence of S, M, and E proteins of SARS-CoV-2, MAVS which is a key adaptor protein in the innate immune response and a critical component in the activation of the type I interferon pathway, was downregulated. MAVS is responsible for sensing viral components and initiating downstream signalling that leads to interferon production ^[69]^. The down-regulation of MAVS is likely to affect NF-κB signalling, as MAVS is involved in the activation of NF-κB during viral infection ^[69,70]^. Consequently, the decreased MAVS expression may result in impaired NF-κB activation. Similarly, SP1, a transcription factor involved in the regulation of various genes, including those associated with immune responses, was also downregulated. Despite the down-regulation of MAVS and SP1, the observed up-regulation of interferon in the transcriptome data suggests the involvement of compensatory or alternative mechanisms in the regulation of interferon response during SARS-CoV-2 protein transfection.

Based on the GO analysis, a significant enrichment of up-regulated genes in the presence of SARS-CoV-2 proteins was observed in biological processes. Specifically, these upregulated genes were found to be enriched in ER-related processes, cellular response to interferon type I, and defence response to viruses. Reactome pathway analysis further supported these findings, revealing similar enriched pathways among the upregulated genes in the presence of SARS-CoV-2 proteins. These pathways included cytokine signalling in the immune system, interferon alpha and beta signalling, antiviral mechanisms stimulated by interferons, unfolded protein response, regulation of HSF1-mediated heat shock response, and activation of IRE1-alpha and chaperones. The funding is consistent with a study by Sun et al. investigating the transcriptome of SARS-CoV-2-infected cells ^[63]^. The SARS-CoV-2 S, M, and E protein datasets exhibited similar patterns of pathway activation, suggesting a common impact on cellular processes. However, the SARS-CoV-2 N protein dataset displayed distinct differences, with the activation of additional pathways involved in immune signalling, such as TRAF6-mediated IRF7 activation and TRAF6-mediated NF-κB activation. Interestingly, pathways associated with ER stress did not show significant enrichment in the N protein dataset.

Among the downregulated genes, all datasets shared common terms for BPs the common DGEs were enriched in viral transcription, translational initiation, non-coding RNA metabolism and ribosome biogenesis and ion transport regulation. The decreased expression of ribosomal proteins in the context of viral protein transfection could indicate a disruption or inhibition of the host’s translational machinery. While increased expression of ribosomal proteins is commonly associated with enhanced protein synthesis, the decreased expression suggests that the viral proteins may interfere with normal ribosome function or protein translation processes ^[71,72]^. This disruption could be a result of the viral proteins manipulating cellular mechanisms, such as inhibiting ribosome assembly or inducing specific modifications to ribosomal proteins. The Reactome analysis revealed consistent pathway suppression across multiple SARS-CoV-2 protein datasets. The NOTCH1 signalling pathway was found to be suppressed in the S, M, E and N protein datasets. Suppression of ion channel transport was observed in the S and E protein datasets. Additionally, the M and E protein datasets exhibited pathway suppression in the context of ERBB4 signalling. Further found to be supressed in N dataset includes influenza infection, G alpha 12/13 signalling events, SEMA4D signalling, RAB protein-mediated trafficking, and muscle contraction. Among the common pathway suppressions observed across multiple SARS-CoV-2 protein datasets, the NOTCH1 signalling pathway and ion channel transport appear to be consistent. Notch signalling has been linked to several biological processes mediating viral infections ^[73,74]^. One study found that proteins interacting with SARS-CoV-2 RNA were associated with Notch2 receptor signalling, suggesting a potential connection between the virus and the Notch pathway ^[75]^. The suppression of the Notch pathway may indicate a disruption of its normal functions, potentially impacting its regulatory role in the context of viral infection.

There have been several studies reporting dysregulation of ion transport in SARS-CoV-2-infected cells ^[76–78]^ which aligns with our findings of downregulated ion transport pathways in the presence of SARS-CoV-2 proteins. The down-regulation of ion transport observed in our transcriptome data, along with the activation of ER stress pathways, is consistent with previous studies that have reported decreased mRNA and protein levels of Na,K-ATPase subunits in SARS-CoV-2-infected cells and postmortem lung tissue samples from COVID-19 patients ^[78,77]^. These findings suggest a potential association between SARS-CoV-2 infection, ER stress, and impaired ion transport. The impairment of Na,K-ATPase maturation and its delivery to the cell membrane, resulting from disrupted chaperone-assisted folding in the ER lumen, as well as the possible interference of the SARS-CoV-2 spike protein with glycosylation-dependent protein folding machinery, may contribute to the observed decrease in Na,K-ATPase abundance at the plasma membrane ^[78,77]^. Together, these findings highlight the intricate interplay between viral infection, cellular processes, and dysregulation of ion transport, providing valuable insights into the pathogenesis of COVID-19.

In conclusion, our comprehensive comparative analysis of individual SARS-CoV-2 structural proteins has revealed compelling insights into the molecular mechanisms driving COVID-19 pathogenesis. The findings demonstrate the potential impact of these viral proteins on host immune responses, transcriptional regulation, and cellular stress pathways. We observed robust up-regulation of interferon-stimulated genes (ISGs) and activation of antiviral defence pathways. The presence of SARS-CoV-2 structural proteins induced the up-regulation of ISGs, highlighting their role in modulating host immune defences. Additionally, the activation of HSPs and the UPR suggested the imposition of ER stress by SARS-CoV-2 proteins. Notably, the N protein exhibited distinct regulatory properties, with less stimulation of HSPs compared to other viral proteins. This finding supports the findings that the down-modulation of HIV-1 LTR activation was associated with the up-regulation of ER stress responses.

Dysregulation of transcription factors indicated their involvement in the cellular response to SARS-CoV-2 infection. The differential effects of the N protein on transcription factors highlight its unique role in modulating gene expression patterns. These findings underscore the complexity of SARS-CoV-2-host interactions and the need for further research to understand the underlying molecular mechanisms. Advancing our understanding of these interactions will be crucial for developing targeted therapeutic approaches for COVID-19. In summary, our study provides compelling evidence of the intricate interplay between SARS-CoV-2 structural proteins, host immune responses, transcriptional regulation, and cellular stress pathways, shedding light on COVID-19 pathogenesis.

## Acknowledgements.

We would also like to thank Prof. Andrew Owens for providing the HEK293T/ACE-2 cells and Prof. Paul McNamara for providing the BEAS-2B cells. We would also like to thank Dr Mark Whitehead for assistance with confirming our bioinformatic analyses.

## Author Contributions

WA, JT, GP and WAP conceived the concept and the experiments. WA, FM, KJR, AA, AK conducted the experiments. WA, JT, GP and WAP analysed results. WA wrote the initial manuscript and all authors contributed to writing the final article.

## Data availability

The authors welcome requests for access to the data generated in this study which will be made available by the corresponding author upon reasonable request.

## Additional information

The authors declare no competing interests.

## REFERENCES

1. Chan, J.F.-W., et al., Genomic characterization of the 2019 novel human-pathogenic coronavirus isolated from a patient with atypical pneumonia after visiting Wuhan. Emerging microbes & infections, 2020. 9(1):221–36

2. Bosch, B. J., Van der Zee, R., De Haan, C. A., & Rottier, P. J. (2003). The coronavirus spike protein is a class I virus fusion protein: structural and functional characterization of the fusion core complex. Journal of virology, 77(16), 8801–8811.

3. Hoffmann, M., Kleine-Weber, H., Schroeder, S., Krüger, N., Herrler, T., Erichsen, S., Schiergens, T. S., Herrler, G., Wu, N.-H., & Nitsche, A. (2020). SARS-CoV-2 cell entry depends on ACE2 and TMPRSS2 and is blocked by a clinically proven protease inhibitor. Cell, 181(2), 271–280. e278.

4. Walls, A. C., Park, Y.-J., Tortorici, M. A., Wall, A., McGuire, A. T., & Veesler, D. (2020). Structure, function, and antigenicity of the SARS-CoV-2 spike glycoprotein. Cell, 181(2), 281–292. e286.

5. Wang, Q., Zhang, Y., Wu, L., Niu, S., Song, C., Zhang, Z., Lu, G., Qiao, C., Hu, Y., & Yuen, K.-Y. (2020b). Structural and functional basis of SARS-CoV-2 entry by using human ACE2. Cell, 181(4), 894–904. e899.

6. Magazine, N., Zhang, T., Wu, Y., McGee, M. C., Veggiani, G., & Huang, W. (2022). Mutations and evolution of the SARS-CoV-2 spike protein. Viruses, 14(3), 640.7.

7. de Haan, C. A., & Rottier, P. J. (2005). Molecular interactions in the assembly of coronaviruses. Advances in virus research, 64, 165–230.

8. Stertz, S., Reichelt, M., Spiegel, M., Kuri, T., Martínez-Sobrido, L., García-Sastre, A., Weber, F., & Kochs, G. (2007). The intracellular sites of early replication and budding of SARS-coronavirus. Virology, 361(2), 304–315.

9. Perrier, A., Bonnin, A., Desmarets, L., Danneels, A., Goffard, A., Rouillé, Y., Dubuisson, J., & Belouzard, S. (2019). The C-terminal domain of the MERS coronavirus M protein contains a trans-Golgi network localization signal. Journal of Biological Chemistry, 294(39), 14406–14421.

10. Klein, S., Cortese, M., Winter, S. L., Wachsmuth-Melm, M., Neufeldt, C. J., Cerikan, B., Stanifer, M. L., Boulant, S., Bartenschlager, R., & Chlanda, P. (2020). SARS-CoV-2 structure and replication characterized by in situ cryo-electron tomography. Nature communications, 11(1), 5885.

11. Ron, D., & Walter, P. (2007). Signal integration in the endoplasmic reticulum unfolded protein response. Nature reviews Molecular cell biology, 8(7), 519–529.

12. Hetz, C. and F.R. Papa, The unfolded protein response and cell fate control. Molecular cell, 2018. 69(2):169–81

13. Janssens, S., Pulendran, B., & Lambrecht, B. N. (2014). Emerging functions of the unfolded protein response in immunity. Nature immunology, 15(10), 910–919.

14. Chen, Y., Liu, Q., & Guo, D. (2020). Emerging coronaviruses: genome structure, replication, and pathogenesis. Journal of medical virology, 92(4), 418–423.

15. Huang, C., Wang, Y., Li, X., Ren, L., Zhao, J., Hu, Y., Zhang, L., Fan, G., Xu, J., & Gu, X. (2020a). Clinical features of patients infected with 2019 novel coronavirus in Wuhan, China. The lancet, 395(10223), 497–506.

16. Lucas, C., Wong, P., Klein, J., Castro, T. B., Silva, J., Sundaram, M., Ellingson, M. K., Mao, T., Oh, J. E., & Israelow, B. (2020). Longitudinal analyses reveal immunological misfiring in severe COVID-19. Nature, 584(7821), 463–469.

17. Hojyo, S., Uchida, M., Tanaka, K., Hasebe, R., Tanaka, Y., Murakami, M., & Hirano, T. (2020). How COVID-19 induces cytokine storm with high mortality. Inflammation and regeneration, 40, 1–7.

18. Kedzierska, K., & Crowe, S. M. (2001). Cytokines and HIV-1: interactions and clinical implications. antiviral Chemistry and Chemotherapy, 12(3), 133–150.

19. Huang, K. J., Su, I. J., Theron, M., Wu, Y. C., Lai, S. K., Liu, C. C., & Lei, H. Y. (2005). An interferon-γ-related cytokine storm in SARS patients. Journal of medical virology, 75(2), 185–194.

20. Haque, A., Hober, D., & Kasper, L. H. (2007). Confronting potential influenza A (H5N1) pandemic with better vaccines. Emerging infectious diseases, 13(10), 1512.

21. Sadeghi, M., Eckerle, I., Daniel, V., Burkhardt, U., Opelz, G., & Schnitzler, P. (2011). Cytokine expression during early and late phase of acute Puumala hantavirus infection. BMC immunology, 12(1), 1–10.

22. Sun, S.-C. and S.C. Ley (2008) New insights into NF-κB regulation and function. Trends in immunology. 29(10):469–78

23. Perkins, N.D. (2007) Integrating cell-signalling pathways with NF-κB and IKK function. Nature reviews Molecular cell biology. 8(1):49–62.

24. Chen, L.-F., & Greene, W. C. (2004). Shaping the nuclear action of NF-κB. Nature reviews Molecular cell biology, 5(5), 392–401.

25. Zhang, H. and S.-C. Sun. (2015) NF-κB in inflammation and renal diseases. Cell & bioscience.5(1):1–12

26. Kagoya, Y., Yoshimi, A., Kataoka, K., Nakagawa, M., Kumano, K., Arai, S., Kobayashi, H., Saito, T., Iwakura, Y., & Kurokawa, M. (2014). Positive feedback between NF-κB and TNF-α promotes leukemia-initiating cell capacity. The Journal of clinical investigation, 124(2), 528–542.

27. Rahman, M. M., & McFadden, G. (2011). Modulation of NF-κB signalling by microbial pathogens. Nature Reviews Microbiology, 9(4), 291–306.

28. Neufeldt, C.J., et al., SARS-CoV-2 infection induces a pro-inflammatory cytokine response through cGAS-STING and NF-κB. Communications biology, 2022. 5(1):45

29. Khan, S. Z., Gasperino, S., & Zeichner, S. L. (2019). Nuclear Transit and HIV LTR Binding of NF-κB Subunits Held by IκB Proteins: Implications for HIV-1 Activation. Viruses, 11(12), 1162.

30. Nabel, G., & Baltimore, D. (1987). An inducible transcription factor activates expression of human immunodeficiency virus in T cells. Nature, 326(6114), 711–713.

31. Perkins, N.D., et al., A cooperative interaction between NF-kappa B and Sp1 is required for HIV-1 enhancer activation. The EMBO journal, 1993. 12(9):3551–3558

32. Basseres, D., & Baldwin, A. (2006). Nuclear factor-κB and inhibitor of κB kinase pathways in oncogenic initiation and progression. Oncogene, 25(51), 6817–6830.

33. Hayden, M. S., & Ghosh, S. (2004). Signaling to NF-κB. Genes & development, 18(18), 2195–2224.

34. McKay, L. G., Thomas, J., Albalawi, W., Fattaccioli, A., Dieu, M., Ruggiero, A., McKeating, J. A., Ball, J. K., Tarr, A. W., & Renard, P. (2022). The HCV Envelope Glycoprotein Down-Modulates NF-κB Signalling and Associates With Stimulation of the Host Endoplasmic Reticulum Stress Pathway. Frontiers in Immunology, 13.

35. Zufferey, R., et al. (1997) Multiply attenuated lentiviral vector achieves efficient gene delivery in vivo. Nature biotechnology. 15(9):871–875.

36. Wood, D.E., J. Lu, and B. Langmead (2019) Improved metagenomic analysis with Kraken 2. Genome biology. 20:1–13.

37. Li, H., (2018) Minimap2: pairwise alignment for nucleotide sequences. Bioinformatics. 34(18):3094–3100

38. Law, C.W., et al. (2014) voom: Precision weights unlock linear model analysis tools for RNA-seq read counts. Genome biology,. 15(2):1–17.

39. Ritchie, M. E., Phipson, B., Wu, D., Hu, Y., Law, C. W., Shi, W., & Smyth, G. K. (2015). limma powers differential expression analyses for RNA-sequencing and microarray studies. Nucleic acids research, 43(7), e47–e47.

40. Yu, G., Wang, L.-G., Han, Y., & He, Q.-Y. (2012). clusterProfiler: an R package for comparing biological themes among gene clusters. Omics: a journal of integrative biology, 16(5), 284–287.

41. Subramanian, A., Tamayo, P., Mootha, V. K., Mukherjee, S., Ebert, B. L., Gillette, M. A., Paulovich, A., Pomeroy, S. L., Golub, T. R., & Lander, E. S. (2005). Gene set enrichment analysis: a knowledge-based approach for interpreting genome-wide expression profiles. Proceedings of the National Academy of Sciences, 102(43), 15545–15550.

42. Luo, W., & Brouwer, C. (2013). Pathview: an R/Bioconductor package for pathway-based data integration and visualization. bioinformatics, 29(14), 1830–1831.

43. Stertz, S., Reichelt, M., Spiegel, M., Kuri, T., Martínez-Sobrido, L., García-Sastre, A., Weber, F., & Kochs, G. (2007). The intracellular sites of early replication and budding of SARS-coronavirus. Virology, 361(2), 304–315.

44. de Haan, C. A., & Rottier, P. J. (2005). Molecular interactions in the assembly of coronaviruses. Advances in virus research, 64, 165–230.

45. Chan, C.P., Siu, K.L., Chin, K.T., Yuen, K.Y., Zheng, B. and Jin, D.Y., 2006. Modulation of the unfolded protein response by the severe acute respiratory syndrome coronavirus spike protein. Journal of virology, 80(18), pp.9279–9287.

46. Rosa-Fernandes, L., Lazari, L.C., da Silva, J.M., de Morais Gomes, V., Machado, R.R.G., dos Santos, A.F., Araujo, D.B., Coutinho, J.V.P., Arini, G.S., Angeli, C.B. and de Souza, E.E., 2021. SARS-CoV-2 activates ER stress and Unfolded protein response. BioRxiv, pp.2021–06.

47. McBride, R., Van Zyl, M. and Fielding, B.C., 2014. The coronavirus nucleocapsid is a multifunctional protein. Viruses, 6(8), pp.2991–3018.

48. V’kovski, P., Kratzel, A., Steiner, S., Stalder, H. and Thiel, V., 2021. Coronavirus biology and replication: implications for SARS-CoV-2. Nature Reviews Microbiology, 19(3), pp.155–170.

49. MacMicking, J. D. (2012). Interferon-inducible effector mechanisms in cell-autonomous immunity. Nature Reviews Immunology, 12(5), 367–382.

50. Uematsu, S., & Akira, S. (2006). Innate immune recognition of viral infection. Uirusu, 56(1), 1–8.

51. Yin, X., Riva, L., Pu, Y., Martin-Sancho, L., Kanamune, J., Yamamoto, Y., Sakai, K., Gotoh, S., Miorin, L., & De Jesus, P. D. (2021). MDA5 governs the innate immune response to SARS-CoV-2 in lung epithelial cells. Cell reports, 34(2), 108628.

52. Rebendenne, A., Chaves Valadão, A. L., Tauziet, M., Maarifi, G., Bonaventure, B., McKellar, J., Planès, R., Nisole, S., Arnaud-Arnould, M., & Moncorgé, O. (2021). SARS-CoV-2 triggers an MDA-5-dependent interferon response which is unable to control replication in lung epithelial cells. Journal of virology, 95(8), e02415–02420.

53. Onodi, F., Bonnet-Madin, L., Meertens, L., Karpf, L., Poirot, J., Zhang, S.-Y., Picard, C., Puel, A., Jouanguy, E., & Zhang, Q. (2021). SARS-CoV-2 induces human plasmacytoid predendritic cell diversification via UNC93B and IRAK4. Journal of Experimental Medicine, 218(4).

54. Stanifer, M. L., Kee, C., Cortese, M., Zumaran, C. M., Triana, S., Mukenhirn, M., Kraeusslich, H.-G., Alexandrov, T., Bartenschlager, R., & Boulant, S. (2020). Critical role of type III interferon in controlling SARS-CoV-2 infection in human intestinal epithelial cells. Cell reports, 32(1), 107863.

55. Cheemarla, N. R., Watkins, T. A., Mihaylova, V. T., Wang, B., Zhao, D., Wang, G., Landry, M. L., & Foxman, E. F. (2021). Dynamic innate immune response determines susceptibility to SARS-CoV-2 infection and early replication kinetics. Journal of Experimental Medicine, 218(8).

56. Hadjadj, J., Yatim, N., Barnabei, L., Corneau, A., Boussier, J., Smith, N., Péré, H., Charbit, B., Bondet, V., & Chenevier-Gobeaux, C. (2020). Impaired type I interferon activity and inflammatory responses in severe COVID-19 patients. Science, 369(6504), 718–724.

57. Shemesh, M., Aktepe, T. E., Deerain, J. M., McAuley, J. L., Audsley, M. D., David, C. T., Purcell, D. F., Urin, V., Hartmann, R., & Moseley, G. W. (2021). SARS-CoV-2 suppresses IFNβ production mediated by NSP1, 5, 6, 15, ORF6 and ORF7b but does not suppress the effects of added interferon. PLoS pathogens, 17(8), e1009800.

58. Lei, X., Dong, X., Ma, R., Wang, W., Xiao, X., Tian, Z., Wang, C., Wang, Y., Li, L., & Ren, L. (2020). Activation and evasion of type I interferon responses by SARS-CoV-2. Nature communications, 11(1), 3810.

59. Chang, J. J.-Y., Gleeson, J., Rawlinson, D., De Paoli-Iseppi, R., Zhou, C., Mordant, F. L., Londrigan, S. L., Clark, M. B., Subbarao, K., & Stinear, T. P. (2022). Long-read RNA sequencing identifies polyadenylation elongation and differential transcript usage of host transcripts during SARS-CoV-2 in vitro infection. Frontiers in Immunology, 13.

60. Heck, T. G., Ludwig, M. S., Frizzo, M. N., Rasia-Filho, A. A., & Homem de Bittencourt Jr, P. I. (2020). Suppressed anti-inflammatory heat shock response in high-risk COVID-19 patients: lessons from basic research (inclusive bats), light on conceivable therapies. Clinical Science, 134(15), 1991–2017.

61. Zhang, X., & Yu, W. (2022). Heat shock proteins and viral infection. Frontiers in Immunology, 13.

62. Oglesbee, M. J., Pratt, M., & Carsillo, T. (2002). Role for heat shock proteins in the immune response to measles virus infection. Viral Immunology, 15(3), 399–416.

63. Sun, G., Cui, Q., Garcia Jr, G., Wang, C., Zhang, M., Arumugaswami, V., Riggs, A. D., & Shi, Y. (2021). Comparative transcriptomic analysis of SARS-CoV-2 infected cell model systems reveals differential innate immune responses. Scientific Reports, 11(1), 17146.

64. Satpathy, A. T., & Chang, H. Y. (2015). Long noncoding RNA in hematopoiesis and immunity. Immunity, 42(5), 792–804.

65. Atianand, M. K., Caffrey, D. R., & Fitzgerald, K. A. (2017). Immunobiology of long noncoding RNAs. Annual review of immunology, 35, 177–198.

66. Liu, W., Wang, Z., Liu, L., Yang, Z., Liu, S., Ma, Z., Liu, Y., Ma, Y., Zhang, L., & Zhang, X. (2020). LncRNA Malat1 inhibition of TDP43 cleavage suppresses IRF3-initiated antiviral innate immunity. Proceedings of the National Academy of Sciences, 117(38), 23695–23706.

67. Shaath, H., Vishnubalaji, R., Elkord, E., & Alajez, N. M. (2020). Single-cell transcriptome analysis highlights a role for neutrophils and inflammatory macrophages in the pathogenesis of severe COVID-19. Cells, 9(11), 2374.

68. Moazzam-Jazi, M., Lanjanian, H., Maleknia, S., Hedayati, M., & Daneshpour, M. S. (2021). Interplay between SARS-CoV-2 and human long non-coding RNAs. Journal of cellular and molecular medicine, 25(12), 5823–5827.

69. Seth, R. B., Sun, L., Ea, C.-K., & Chen, Z. J. (2005). Identification and characterization of MAVS, a mitochondrial antiviral signaling protein that activates NF-κB and IRF3. Cell, 122(5), 669–682.

70. Lazear, H. M., Lancaster, A., Wilkins, C., Suthar, M. S., Huang, A., Vick, S. C., Clepper, L., Thackray, L., Brassil, M. M., & Virgin, H. W. (2013). IRF-3, IRF-5, and IRF-7 coordinately regulate the type I IFN response in myeloid dendritic cells downstream of MAVS signaling. PLoS pathogens, 9(1), e1003118.

71. Meyuhas, O. (2008). Physiological roles of ribosomal protein S6: one of its kind. International review of cell and molecular biology, 268, 1–37.

72. Hertz, M. I., Landry, D. M., Willis, A. E., Luo, G., & Thompson, S. R. (2013). Ribosomal protein S25 dependency reveals a common mechanism for diverse internal ribosome entry sites and ribosome shunting. Molecular and cellular biology, 33(5), 1016–1026.

73. Aster, J. C. (2014). In brief: Notch signalling in health and disease. The Journal of pathology, 232(1), 1–3.

74. Shang, Y., Smith, S., & Hu, X. (2016). Role of Notch signaling in regulating innate immunity and inflammation in health and disease. Protein & cell, 7(3), 159–174.

75. Vandelli, A., Monti, M., Milanetti, E., Armaos, A., Rupert, J., Zacco, E., Bechara, E., Delli Ponti, R., & Tartaglia, G. G. (2020). Structural analysis of SARS-CoV-2 genome and predictions of the human interactome. Nucleic acids research, 48(20), 11270–11283.

76. Blanco-Melo, D., Nilsson-Payant, B. E., Liu, W.-C., Uhl, S., Hoagland, D., Møller, R., Jordan, T. X., Oishi, K., Panis, M., & Sachs, D. (2020). Imbalanced host response to SARS-CoV-2 drives development of COVID-19. Cell, 181(5), 1036–1045. e1039.

77. Jha, P. K., Vijay, A., Halu, A., Uchida, S., & Aikawa, M. (2021). Gene expression profiling reveals the shared and distinct transcriptional signatures in human lung epithelial cells infected with SARS-CoV-2, MERS-CoV, or SARS-CoV: potential implications in cardiovascular complications of COVID-19. Frontiers in cardiovascular medicine, 7, 623012.

78. Kryvenko, V., & Vadasz, I. (2021). Molecular mechanisms of Na, K-ATPase dysregulation driving alveolar epithelial barrier failure in severe COVID-19. 320(6), L1186–L1193.

